# Co-chaperone BAG3 directly target autophagic degradation via its LC3-interacting regions

**DOI:** 10.1101/2023.02.01.526551

**Authors:** Hagen Körschgen, Marius Baeken, Daniel Schmitt, Heike Nagel, Christian Behl

## Abstract

The co-chaperone BAG3 is a hub for a variety of cellular pathways via its multiple domains and its interaction with HSP70 and HSPB8. Under aging and cellular stress conditions in particular, together with molecular chaperones, BAG3 ensures the sequestration of aggregated or aggregation prone ubiquitinated proteins to the autophagic-lysosomal system via ubiquitin receptors. There are emerging indications that BAG3-mediated selective macroautophagy also copes with non-ubiquitinated cargo. Phylogenetically, BAG3 comprises several highly conserved predicted LIRs, LC3-interacting regions, which might directly target BAG3 including its cargo to ATG8 proteins and directly drive their autophagic degradation. Based on pull-down experiments, peptide arrays and proximity ligation assays, our results provide evidence of an interaction of BAG3 with ATG8 proteins. In addition, we could demonstrate that mutations within the LIRs impair co-localization with ATG8 proteins in immunofluorescence. A BAG3 variant mutated in all LIRs results in a substantial decrease of BAG3 levels within purified native autophagic vesicles compared to wild-type BAG3. These results strongly suggest LC3-mediated sequestration of BAG3. Therefore, we conclude that in addition of being a key co-chaperone to HSP70, BAG3 may also act as cargo receptor for client proteins, which would significantly extend the role of BAG3 in selective macroautophagy and protein quality control.

**Synopsis:** BAG3 ensures sequestration of aggregated ubiquitinated proteins to the autophagic-lysosomal degradation. Based on emerging indications this BAG3-mediated macroautophagy may also cope with non-ubiquitinated clients and comprises conserved predicted LC3 interacting regions, we analyzed the interaction with LC3 proteins. We evidenced an interaction of BAG3 with LC3 proteins by various measures including pull-down experiments, peptide arrays, proximity ligation assays, co-localization and native autophagic vesicles analysis. These results suggest BAG3 may additionally act as cargo receptor for client proteins.

**Abstract Figure:** 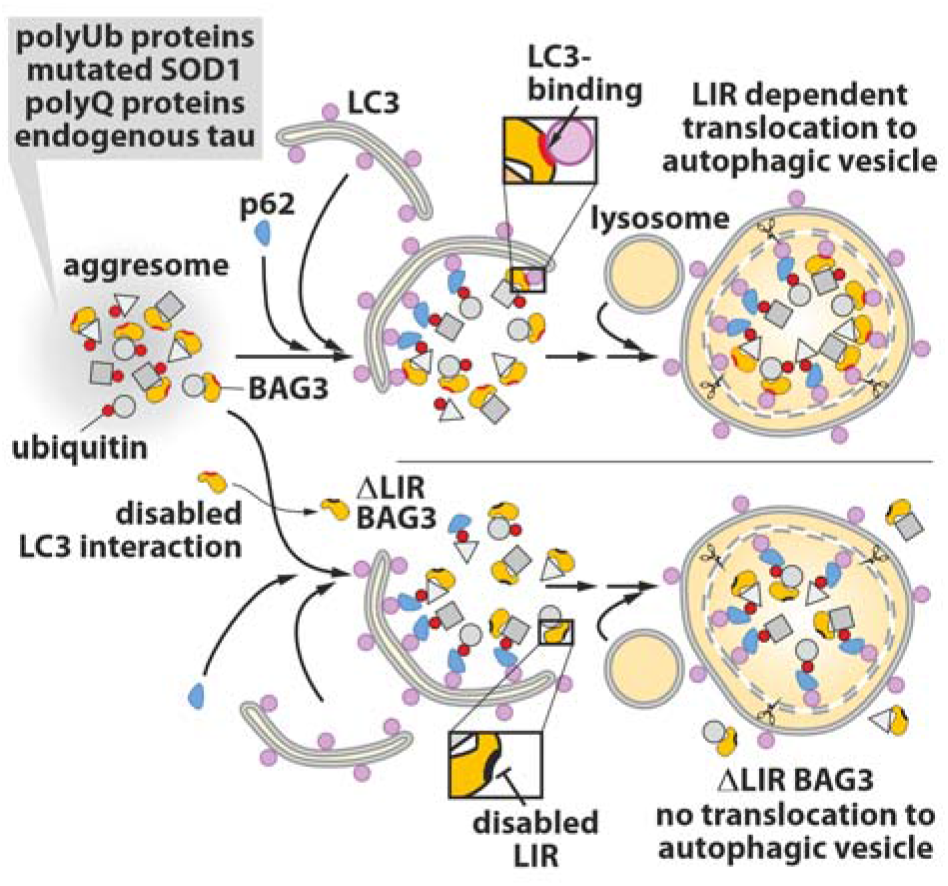

## Introduction

Maintaining proteostasis, particularly under cellular stress conditions that promote protein misfolding or degradation, is crucial for a functional proteome. Accordingly, sophisticated protein quality control mechanisms have evolved, and eukaryotic cells govern the disposal of dysfunctional proteins via the ubiquitin-proteasome system or autophagosome-lysosome pathway. Consequently, it is essential to target such proteins that are to be degraded.

Molecular chaperones, such as HSPA (HSP70s) or HSPB (small HSPs), are able to identify and bind disordered or hydrophobic regions of unfolded proteins (1, 2). In a coordinated multiprotein complex with other chaperones and co-chaperones, those ligands are either refolded to their native state or, if not achievable, sequestered for degradation (3–5). Here, the cytoplasmic co-chaperone BAG3 (Bcl-2 associated athanogene 3) mediates a translocation to introduce the cargo to the autophagic pathway (6–8). Together with HSP70 the multidomain structure, consisting of the WW, PxxP and BAG domains, as well as its IPV and 14-3-3 motifs, enable BAG3 to serve as a hub for proteotoxicity-induced signaling (9). The cargo of small HSPs and HSP70s is attached to BAG3 via the IPV motifs and the BAG domain, respectively (7, 10, 11).

Canonically, BAG3-mediated selective macroautophagy (referred to as autophagy), requires the recognition of the cargo ubiquitination status (12). Ubiquitinated protein aggregates are recognized by adapters/receptors like Nbr1, OPTN or p62/SQSTM1 and targeted to autophagophores via interaction with lipidated ATG8 proteins (13–15).

In most mammals, six genes encode for members of ATG8 related proteins. Respectively, three genes encode either microtubule-associated proteins 1A/1B light chain 3 (LC3A, LC3B and LC3C) or γ-aminobutyric acid receptor-associated proteins (GABARAP, GABARAPL1 and GABARPL2). Conjugated to phosphatidylethanolamine (PE) these are the only membrane-anchored representatives of ubiquitin-like proteins (16).

Most ATG8 interacting proteins comprise an ATG8-interacting motif (AIM) (resp. LC3 interacting region (LIR) or GABARAP interacting motif (GIM)), which is characterized by an aromatic residue spaced by two amino acids a branched, hydrophobic sidechain ([F/W/Y]_0_XX[L/I/V]_3_) (17). In many cases this core motif is flanked by acidic or polar residues, predominantly N-terminally (18). Accordingly, the phosphorylation of the polar residues in many ligands positively modulates binding to LC3 (19–21).

Additional non-canonical binding motifs, which do lack the aromatic residue interacting with the hydrophobic pocket (HP1) of LC3 proteins, as well as binding via accessory interaction sites have been described recently (22–25). In the past years, few studies addressed the binding preferences and specificity of ATG8s (21, 26, 27). Thus far, some ligand preferences have been characterized (28).

The different binding modes and the varying preferences of the ATG8 proteins illustrate the complexity in regulating the sequestration of autophagic cargo. Apart from the *bona fide* autophagy receptor p62, which links ubiquitinated cargo with ATG8 proteins, there is emerging indication of ubiquitin independent autophagic degradation of cargo sequestrated by BAG3 (8, 29). A direct interaction of BAG3 with ATG8 proteins would decipher this degradation. Indeed, *in silico* analyses identified five potential LIR motifs in BAG3 that are conserved within the craniotes (30). In human BAG3, these include Y^86^ PQL^89^, Y^93^ IPI^96^, Y^205^ ISI^208^, Y^247^ HKI^250^, and Y^451^ LMI^454^.

Therefore, we further investigated BAG3 for interaction with ATG8 proteins. Our results strongly suggest the functionality of these LIRs. Mutations within these resulted in a significant reduction of BAG3 in autophagic vesicles. These findings suggest the novel function of BAG3 as an adapter for non-ubiquitinated cargo in addition to its co-chaperone activity, significantly expanding the role of BAG3 in selective autophagy.

## Materials and Methods

### Antibodies

**Table 1:**
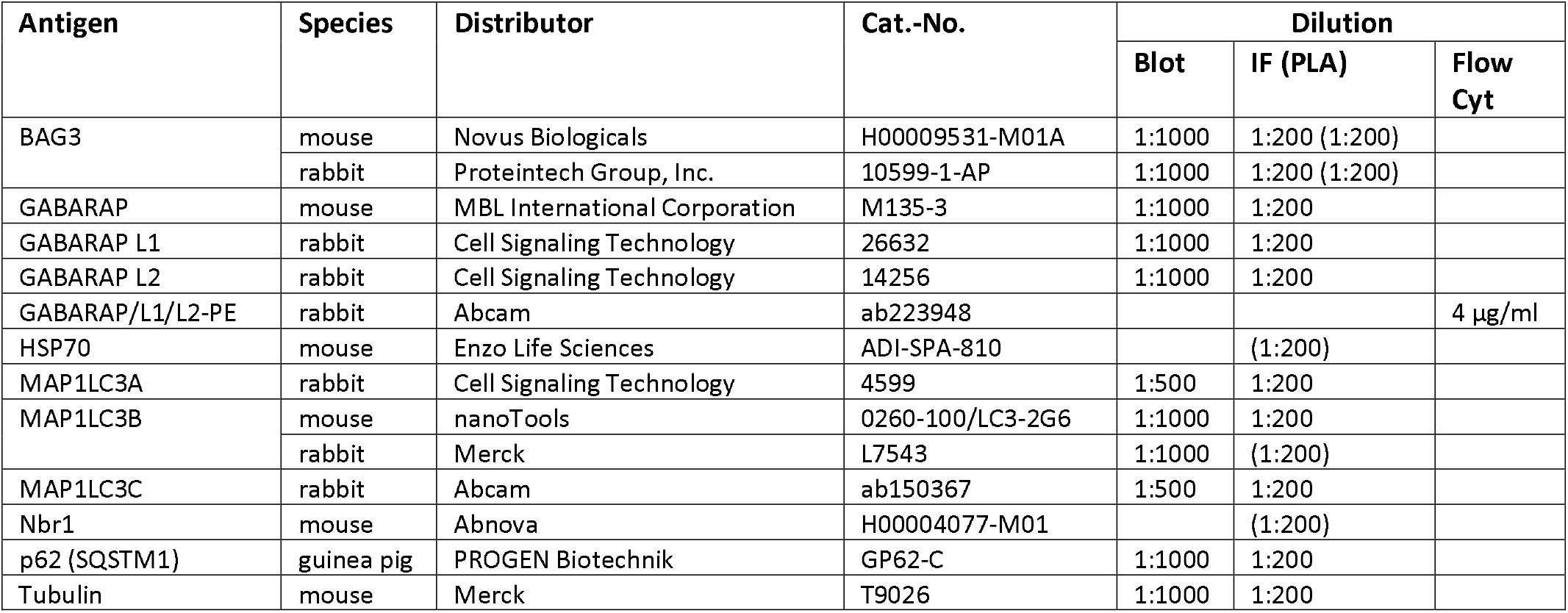
List of all antibodies used

### Cell culture and treatment

HeLa cells (Clone HR5-CL11) (31) were used to establish a BAG3 deficient cell line via CRISPR-Cas9 technology (32) using 5’-GACCGGCTGGCCCTTCTTCG-3’ as BAG3 gRNA. The cells were transfected with three different plasmids each encoding a unique U6 driven gRNA sequence and a CMV driven Cas9 nuclease. 48 h post transfection, the cells were singularized into 96-well plates and the knock-out confirmed by sequencing and immunoblot (Figure 1A). Cells were cultured in Dulbecco’s Modified Eagle Medium (gibco, 41965-039) supplemented with 10% (v/v) fetal bovine serum, 1 mM sodium pyruvate (gibco, 11360039) and 1% antibiotic/antimycotic solution (Merck, A5955). Medium was refreshed every 3 days during cultivation and every 24h in an experimental setting. Cells were treated with stock solutions of MG-132 (Merck, 474790), PYR-41 (Merck, 662105) or Bafilomycin A1 (Biozol, TOR-B110000) in DMSO as specified.

**Figure 1.**
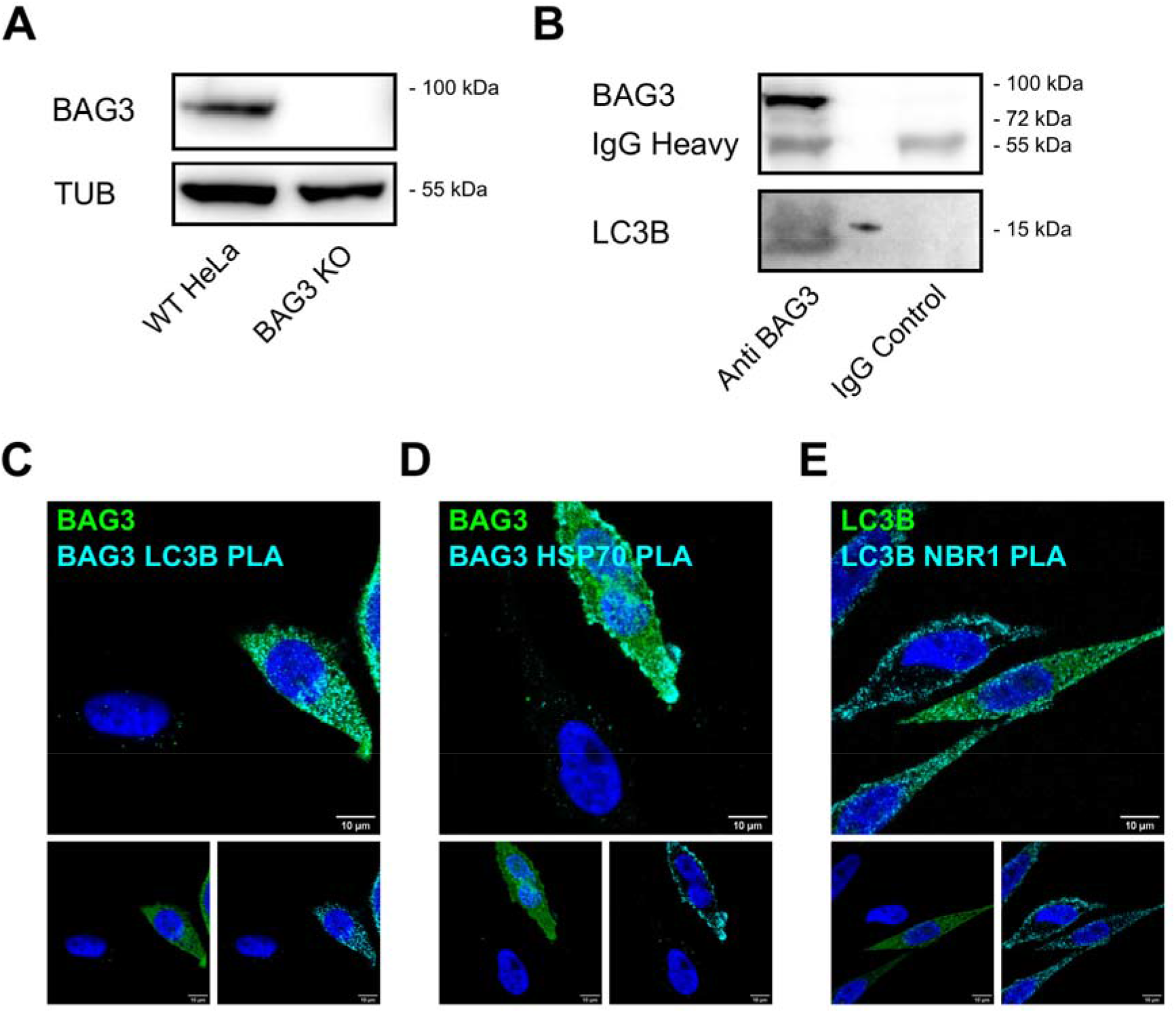
Detection of BAG3-LC3B interaction via immunoprecipitation and proximity ligation assay (PLA). (A) *BAG3*^-/-^ in HeLa cells was confirmed by immunoblot. (B) *BAG3*^-/-^ cells overexpressing WT-BAG3 were treated with PYR-41 (12.5 μM) for 24h and Bafilomycin A1 (2 μM) for 4h; 48 h post transfection whole cell lysates were obtained and immunoprecipitated using an anti-BAG3 antibody or an IgG control.(C-E) PLA for confirmation of BAG3-LC3B interaction (cyan) (C), BAG3-HSP70 interaction (D) and LC3B-Nbr1 interaction (E). All PLA were performed on *BAG3* ^-/-^ cells transiently transfected with human BAG3-eGFP and treated with MG-132 (10 h, 25 μM), Bafilomycin A1 (4 h, 2 μM) and PYR-41 (4 h, 12.5 μM). Representative images obtained from three biological replicates are displayed. Magnification: 100x. Scale bar: 10 μm.

### Autophagic vesicle purification

Autophagic vesicles were purified based on a purification method previously described (33). At least 1 × 10^7^ cells were collected using Trypsin/EDTA and centrifuged at 306 x g for 4 min. After resuspension in PBS supplemented with cOmplete™ EDTA-free (Roche), cell disruption was performed using a UP50H ultrasonic processor (Hielscher) for 3 × 2 s with an amplitude of 60 %. Samples were then centrifuged at 3 000 x g for 10 min at 4 °C, supernatants were collected and centrifuged at 18 620 x g for 1 h at 4 °C. Pellets were washed and resuspended in PBS and subsequently incubated with 4 μg/ml of PE-conjugated GABARAP/GABARAPL1/GABARAPL2 antibody (Abcam, ab223948) for 1 h. The samples were centrifuged again with 18 620 x g for 1 h at 4 °C, pellets were washed and resuspended in PBS. For autophagic vesicle sorting a BD FACSAria III SORP (BD Biosciences) equipped with a 70 μm nozzle and a 1.0 FSC neutral density filter was used. The compartment containing autophagic vesicles was first established using an FSC/SSC plot on a logarithmic scale, followed by a doublet discrimination gate using SSC-A/W. Autophagic vesicles were defined as PE-positive events (488 nm, BP 530/30), whose positivity was conducted according to the background given by an unstained negative control. Vesicle sorting was achieved using minimum speed (flow rate < 3.0) maintaining less than 19 000 events per second. Analysis was performed using FlowJo v10.6.1 (BD Biosciences). Proteins of isolated autophagic vesicles were analysed using a methanol/chloroform (2:1) precipitation protocol and subsequent resuspension in urea buffer (8 M urea and 4 % (w/v) CHAPS in 30 mM Tris (pH 8.5 with HCl)), including EDTA-free protease inhibitor. For immunoblot quantification, we analyzed each 2 million autophagic vesicles by normalizing according to the initial amount of tubulin and expression level of BAG3 and ΔLIR, respectively.

### Immunoprecipitation

Cells were lysed in immunoprecipitation buffer (50 mM Tris, pH 7.4, 150mM NaCl, 2 mM Na_2_EDTA, 1 mM Na_4_EGTA, 10% glycerol, 1% Triton X-100, 1 mg/ml cOmplete™ EDTA-free) and passed three times through a 23G needle. Protein concentration was determined using BCA assay (Thermo Fisher Scientific, 23225). The lysate (500 μg) was precleared with Pierce™ Protein A/G magnetic beads (ThermoFisher Scientific, 88802), incubated with 2 μl anti-BAG3 antibody (Proteintech, 10599-1-AP) at 4°C and precipitated with fresh beads. After three washing steps elution was accomplished via reducing SDS loading buffer. Samples were separated via SDS-PAGE and analyzed via immunoblotting.

### Co-localization analysis by immunofluorescence microscopy

BAG3 deficient HeLa cells were cultured on glass cover slips and treated as indicated. Cells were then fixed with 4% formaldehyde in PBS (Carl Roth, P087.5), permeabilized with 0.1% Triton X-100 in PBS and blocked using PBS + 3% (w/v) bovine albumin. Antibodies were incubated as specified in PBS + 1% (w/v) albumin. After incubation with primary antibodies, cells were incubated with cyanine (Cy2, Cy3, Cy5) –conjugated antibodies (Jackson ImmunoResearch). Stained cells were imaged with a Zeiss LSM710 confocal microscope. For evaluation of co-localization z-stacks (63x magnification; ≥5 slices; 0.5 μm optical sections) with approximately 10 cells per image were acquired of at least three randomly chosen optical fields from three independent biological replicates. Co-localization of BAG3 and LC3s was quantified using the ImageJ software in form of the percentage of channel intensity that is above the channel threshold that is co-localized.

### Peptide Array

Spot synthesis for peptide array of human BAG3 (15mer peptides, 3 residues shift per spot, N-terminally acetylated) on cellulose membrane was ordered from Intavis Peptides Services (Tübingen, Germany). Protein binding was performed according to the manufacturer’s specification using heterologously expressed and GST-tagged LC3s (LC3A (Novusbio, H00084557-P01), LC3B (Novoprolabs, 509467), LC3C (Abnova, H00440738-P01) at 2 μg/ml. Binding of the proteins was evaluated using an anti-GST-HRP conjugate (GE Healthcare, RPN1236). For comparative evaluation of affinities, the signal intensities of the respective LIR motif containing peptides were averaged and normalized to the averaged intensity of the most intense motif. In the case of peptides containing two LIR motifs, the signals from those overlapping peptides were excluded from evaluation.

### Plasmids and transfection

BAG3 deficient HeLa cells were transiently transfected via calcium phosphate precipitation. Transfected cells were washed 24 h post transfection and fresh medium was applied. Treatment was administered in such a way that the cells were harvested 48 h after transfection. Expression constructs of wild-type human BAG3 and C-terminal eGFP-tagged BAG3 were used as previously published (6). For generation of single point LIR mutants and ΔLIR BAG3 the QuikChange Lightning Site-Directed Mutagenesis Kit (Agilent, 210518) was utilized according to the manufacturer’s instructions. Mutagenesis was confirmed via sequencing.

**Table 2:**
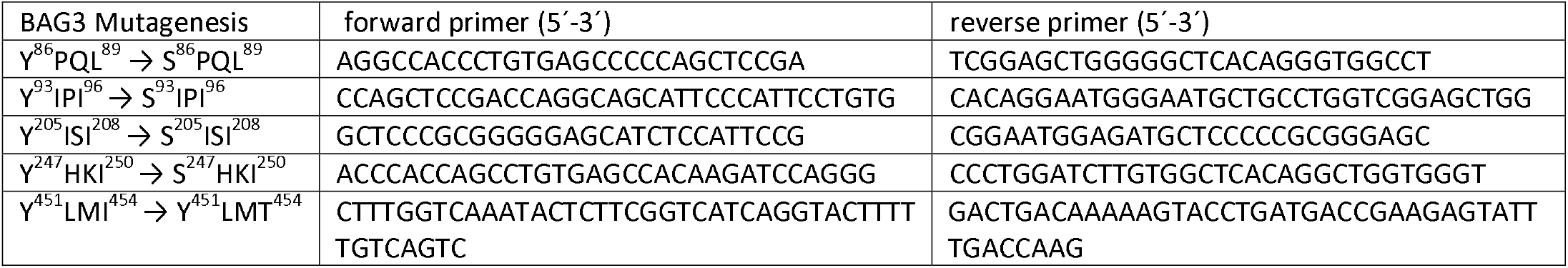
List of primers used for mutagenesis

### Proximity ligation assay (PLA)

*In situ* protein-protein interaction was evidenced using Duolink® Proximity Ligation Assay (Merck, DUO920082, DUO92004, DUO92008). Cells were fixed with formaldehyde and permeabilized using ice-cold 90% methanol. The sample preparation was conducted according to the manufacturer’s protocol with an amplification time of 150 min. Stained cells were imaged with a Zeiss LSM710 confocal microscope.

### Immunoblotting

Reducing SDS-loading buffer was added to the samples, incubated for 5 min at 98 °C and subjected to SDS-PAGE. Gels were then blotted onto nitrocellulose membranes and incubated for 1 h at RT with TBS-T (0.05% Tween-20) with additional 5% non-fat dried milk powder. Antibodies were incubated in TBS-T and detected using peroxidase-conjugated secondary antibodies (Jackson ImmunoResearch) with an Amersham Imager 600 system (GE Scientific).

### Statistical methods

Graphs and statistics were assembled using GraphPad Prism 9 from GraphPad Software, Inc. All statistical analyses were performed according to normal distribution and variance differences by two-way ANOVA followed by original Benjamini and Hochberg posthoc tests. The results display mean ± standard deviation (SD). Statistical significance was accepted at a level of p < 0.05 (p value ≤ 0.05 = *, ≤ 0.01 = **, ≤ 0.001 = ***)

## Results

### BAG3 directly interacts with LC3B

We generated a *BAG3*^*-/-*^ HeLa cell line using a CRISPR/Cas9 approach to prepare a cell line with a defined background (Figure 1A). Then, to confirm the physical functionality of the predicted LIR motifs on protein level, we assessed the interaction via immunoprecipitation of transiently overexpressed wild-type BAG3 and the *bona fide* autophagy marker LC3B. Therefore, the cells were treated with the potent E1 ligase inhibitor Pyr-41 (34) to block ubiquitination dependent protein degradation and allowed accumulation of autophagosomes via bafilomycin A1 (BafA1) to enhance the signal. Hereby, we were able to co-precipitate LC3B with BAG3 (Figure 1B). IgG controls were devoid of BAG3 and LC3B.

Although this *in vitro* pull down is a strong indication of a physiological interaction, it is unclear whether this also occurs within intact cells. Therefore, we performed a proximity ligation assay, which employs immunodetection coupled with an amplification of DNA probes to detect the proximity (max. 40 nm distance) of both proteins. In approximation this method uncovers only direct interactions of two different proteins *in situ*. We were thus able to identify an interaction between BAG3 and LC3B within BAG3-eGFP overexpressing *BAG3* ^*-/-*^ cells (Figure 1C). The negative controls with only the single antibodies against BAG3 and LC3B (Suppl. Figure 1), respectively, as well as the positive controls with well-described interaction partners, for BAG3 (HSP70) and for LC3B (Nbr1), validate the method for detecting the binding of BAG3 to LC3B *in situ* (Figure 1 D & E). The evidence of BAG3-HSP70 interaction is restricted only to cells displaying a BAG3-eGFP expression, whereas the LC3B-Nbr1 interaction is independent of BAG3 expression. Accordingly, detection of the BAG3-LC3B complex, which also occurs only in BAG3-positive cells, is proof of principle for an *in cellulo* interaction.

### BAG3 accumulates in isolated autophagic vesicles

The recent establishment of a purification protocol (33) made it possible to extract and enrich intact native autophagic vesicles from cultured cells by means of a combination of successive centrifugations, an antibody-based fluorescence tagging of ATG8-positive structures and subsequent sorting via flow cytometry (Figure 2A, Suppl. Figure 2). The different centrifugation steps effectively reduced the unlipidated form of LC3B accompanied by enriched levels of lipidated LC3B (Figure 2 B). In accordance with Schmitt et al. the specific detection and sorting of ATG8-positive structures finally resulted in purified vesicles containing prominent autophagic markers such as LC3B and SQSTM1/p62 in the absence of cytosolic proteins (e.g. LC3B-I) or other contaminants like fragments of the cytoskeleton like tubulin (figure 2 B). Here, we treated cells with MG-132 to inhibit proteasomal degradation and to induce the BAG3-mediated selective autophagy as well as with bafilomycin A1 to raise the total quantity of autophagic vesicles. Furthermore, compared to control conditions, we incubated cells with Pyr-41 to diminish the ubiquitylation-dependent degradation pathway (figure 2 C).

**Figure 2.**
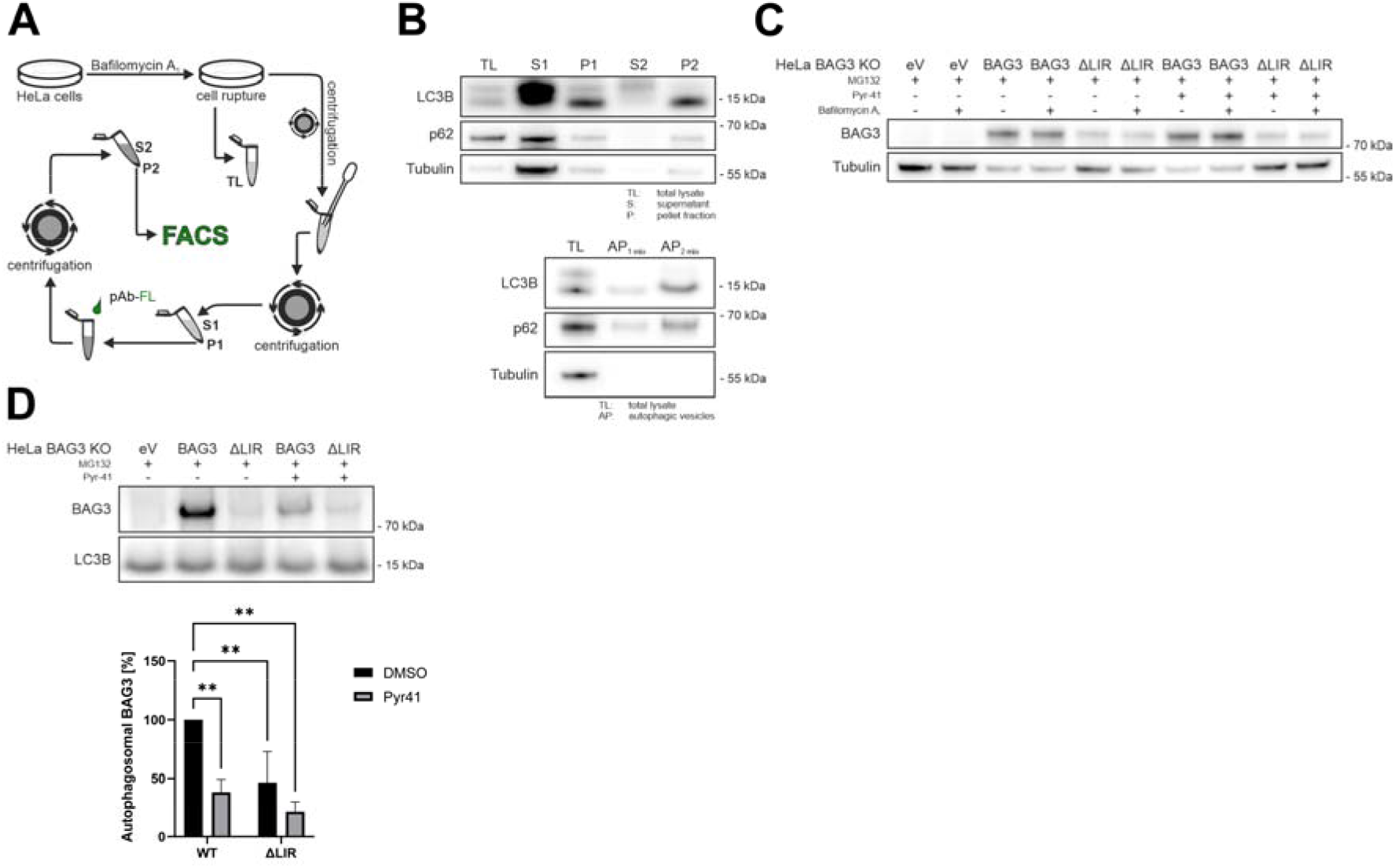
BAG3 in isolated autophagic vesicles. (A) Schematic illustration of autophagic vesicle isolation. (B)Analysis of purification progress by immunoblot; fractions as indicated in A. (C) Immunoblots of total cell lysates from which autophagic vesicles originated. (D) BAG3 detection in autophagic vesicles isolated from *BAG3* ^-/-^ cells overexpressing WT-BAG3 or ΔLIR-BAG3 under indicated treatment paradigms with MG-132 (10 h, 25 μM) and PYR-41 (4 h, 12.5 μM). Bafilomycin A1 (4 h, 2 μM) was always administered. Statistical quantification of autophagosomal BAG3 was performed with 2-way ANOVA followed by a Benjamini Hochberg posthoc test. ‘**’ indicates p<0.01.

For a comparative analysis, whether BAG3 accumulates in autophagosomes in a LIR dependent manner, we created a ΔLIR-BAG3 mutant, which had all potential LIR motifs disabled by changing their tyrosine residue to a serine or, in the case of Y^451^ LMI^454^, the isoleucine residue to a threonine, to reduce possible effects on the integrity of the BAG-domain. *BAG3*^*-/-*^ cells were transfected either with an empty vector (eV), WT-BAG3 or ΔLIR-BAG3. Indeed, this approach confirmed a LIR dependent accumulation of BAG3 in autophagic vesicles, with only minor amounts of ΔLIR-BAG3 present, likely remnants of BAG3 as cargo and not as receptor. Interestingly, additional treatment with Pyr-41 decreased the amount of BAG3. Looking at the total cell lysates we observed a lower expression of ΔLIR-BAG3 compared to wild-type BAG3 (Figure 2D).

### Mutations of potential LIR-motifs within BAG3 abolishes BAG3/LC3 interaction *in cellulo*

We previously reported that other members of the LC3 protein family may compensate a potential loss of LC3B in the autophagic system (35). Therefore, we analyzed whether BAG3 could also interact with LC3A and LC3C, and whether each of these LC3s is selective for a specific potential LIR-motif. To investigate whether BAG3 interacts with LC3A and LC3C we used an immunofluorescence approach and analyzed the co-localization of BAG3 and the LC3s in cells treated with BafA1 and Pyr-41 (Figure 3A, Suppl. Figure 3). We observed a strong correlation between LC3A and BAG3 in BafA1 treated cells independent of presence of Pyr-41. Co-localization of the ΔLIR-BAG3 mutant was significantly reduced (Figure 3B).

**Figure 3.**
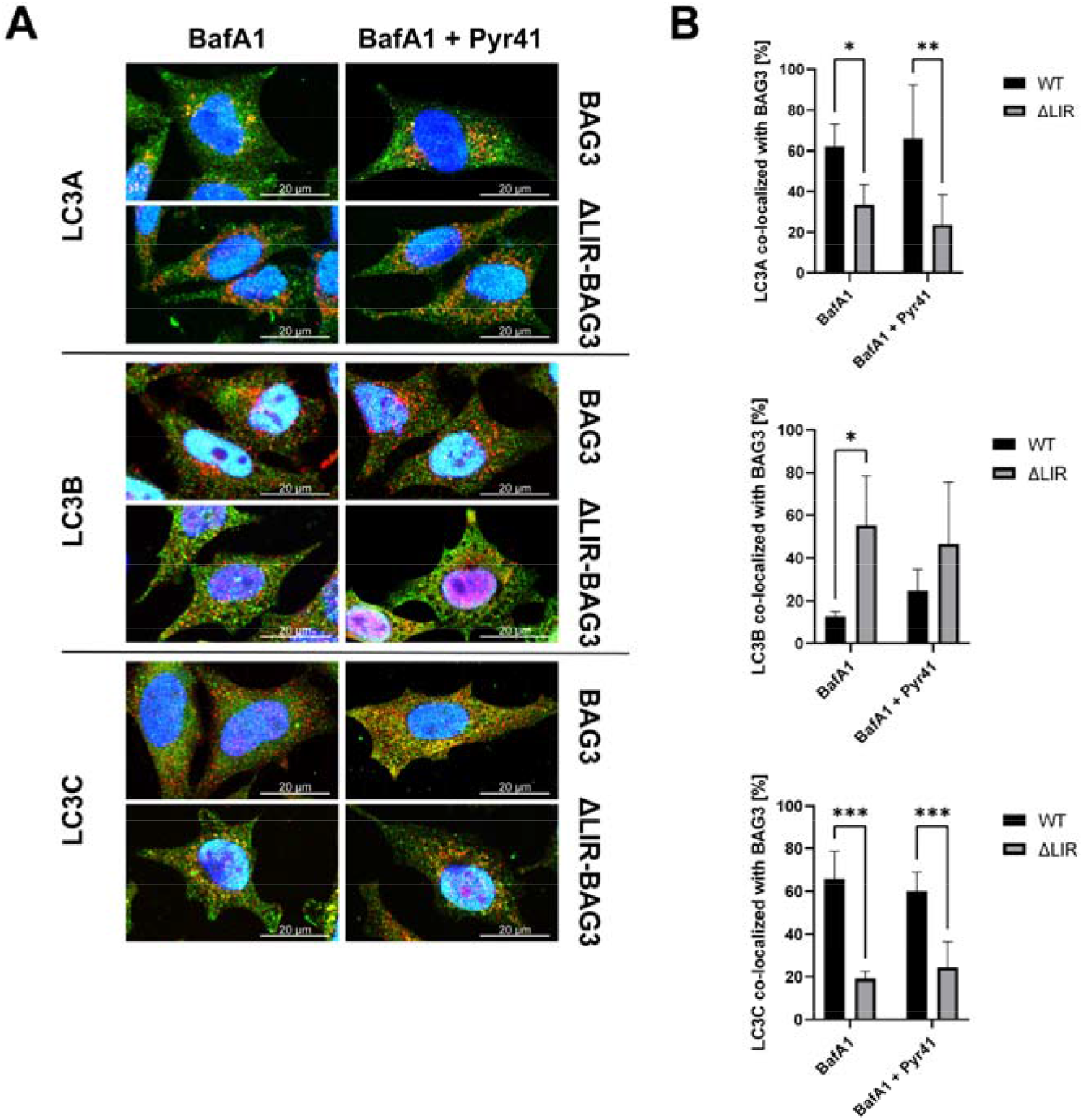
Co-localization analysis of WT and ΔLIR-BAG3 with LC3A, LC3B and LC3C. (A) Representative images of *BAG3* ^-/-^ cells overexpressing WT-BAG3 or ΔLIR-BAG3 treated with Bafilomycin A1 (4 h, 2 μM) and PYR-41 (24 h, 12.5 μM). Magnification: 100x. Scale bar: 20 μm. BAG3 is shown in green, the indicated LC3 in red. (B) Quantification of BAG3 co-localizing with indicated LC3. Statistical significances were reported after 2-way ANOVAs followed by a Benjamini Hochberg posthoc tests. ‘*’ indicates p<0.05, ‘**’ p<0.01 and ‘***’ p<0.001.

Unexpectedly, the co-localization of BAG3 and LC3B was the weakest of all three LC3s. This co-localization, however, was significantly increased for the ΔLIR-BAG3 mutant but was lost with concomitant Pyr-41 treatment. BAG3 features the strongest correlation with LC3C. Like the behavior of LC3A, co-localizations did not change by Pyr-41 treatment. Co-localization was significantly reduced in the ΔLIR-BAG3 mutant for LC3C as for LC3A.

### In depth analysis of putative LIR-motifs in BAG3 reveals LC3 discrepancies

Next, we performed a BAG3 peptide array to assign the binding to the five potential LIRs. Due to the lack of structural information on BAG3 no restriction to particular motifs seemed reasonable. We screened the amino acid sequence for binding regions using synthetic 15mer peptides by shifting each by three residues. The results of the peptide array did not indicate a clear preference for one single putative LIR motif (Figure 4 B, C). For LC3A and LC3B, Y^86^ PQL^89^ containing peptides reveal the highest affinity of the proposed LIRs. The downstream next motif, Y^93^ IPI^96^ displays a much weaker interaction for all LC3s, though a reliable evaluation is problematic due overlap with Y^86^ PQL^89^ peptides. All three LC3 proteins exhibit binding to Y^205^ ISI^208^ -containing peptides, with LC3A demonstrating the strongest intensity. For LC3C, the interaction with Y^247^ HKI^250^ is most intense, although here, as with LC3A^205^ and LC3B^208^, only the peptide containing this sequence at the free N-terminus is recognized. Accordingly, it cannot be concluded that any interaction with this motif is valid based on this approach. Binding to Y^451^ LMI^454^, which is located within the BAG domain mediating the HSP70 interaction, is weak for LC3B and negligible for LC3A and C, respectively. Apart from LC3C whose interaction with the BAG3 peptides is almost exclusively limited to regions containing LIRs, pronounced interactions with peptides without LIR motifs were observed for LC3A and especially for LC3B. Here, based on the recognized sequences, there are currently no data to elucidate whether these are non-canonical binding motifs, as yet unknown interaction mechanisms. Due to high frequency of hydrophobic residues like in the PxxP domain within BAG3, which are also most frequent among the LC3s in LC3B, the signal may likewise be non-specific binding (36).

**Figure 4.**
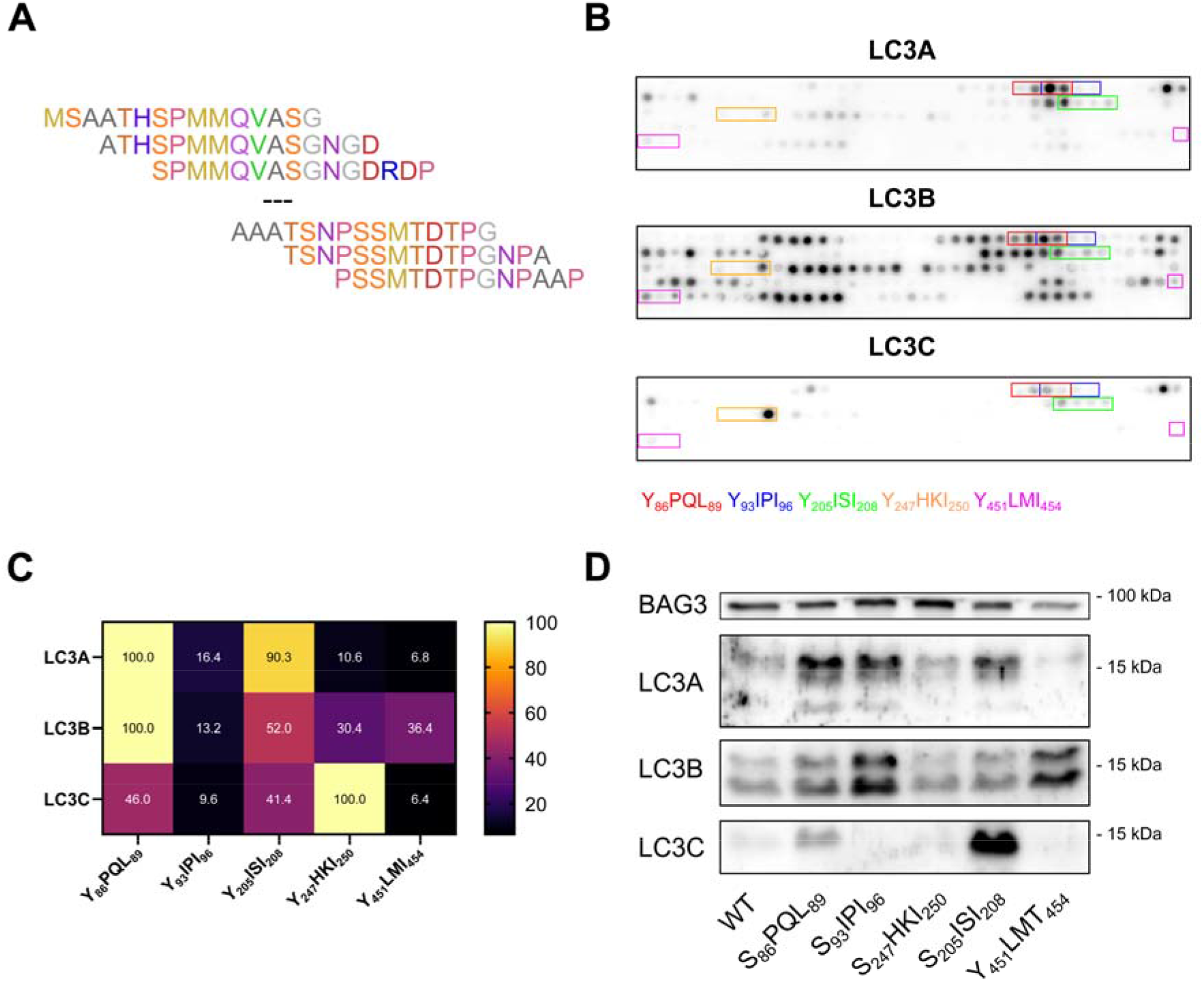
Different LIRs serve different LC3s. (A) Exemplary of the first and last three oligopeptides part of the BAG3 peptide array. With each peptide the primary structure of the oligopeptide shifts by three amino acids towards the C-terminus. (B) Membranes of the BAG3 peptide arrays incubated with the indicated recombinant human GST-tagged LC3 protein. LC3 binding was detected using an anti-GST antibody. Oligopeptides containing the putative LIR motif are highlighted with a unique color for each. (C) Heatmap comparing mean signal intensity of the arrays displayed in B. The motif with the strongest signal intensity was defined as 100%. (D) Immunoprecipitations of *BAG3* ^*-/-*^ cells overexpressing indicated single LIR mutants of BAG3 and detection of co-precipitated LC3s. All cells were treated with Bafilomycin A1 (4 h, 2 μM) and PYR-41 (24 h, 12.5 μM).

In order to analyze the binding properties of each individual putative LIR motif at the protein level, we generated loss-of-function mutations for each motif and tested them for their binding properties via co-immunoprecipitation (figure 4 D). In general, co-immunoprecipitation of LC3B was the highest, while the LC3C interaction appeared weakest. Interestingly, the Y^86^ S mutation led to enhanced binding of all three LC3s, while the Y^93^ S mutation indicated enhanced binding of LC3A and LC3B, but a reduction for LC3C. Mutation of Y^247^ S had limited effects, while Y^205^ S suggested an increase in LC3C binding. Finally, the I^454^ T mutant demonstrated lower binding of LC3A and LC3C, but increased binding to LC3B. None of the single mutants effects an abolishment of LC3B binding. These results match with the results of the peptide array and may reflect compensatory effects of remaining LIRs or LC3 orthologues.

Next, we looked at the co-localization of the single point mutations with LC3A, LC3B and LC3C. For LC3A, within all four treatment groups, we observed a significant increase in co-localization with BAG3 for Y^86^ S, Y^93^ S and Y^205^ S, while the I^454^ T mutant showed a significant decrease (Figure 5 A, B; Suppl. Figure 4 A). These are consistent with the outcome of the co-immunoprecipitation. LC3B, on the other hand demonstrated low levels of co-localization, with the I^454^ T mutant demonstrating a significant increase (Figure 5 C, D; Suppl. Figure 4 B). The increase observed in the co-immunoprecipitations for the Y^93^ S did not yield increased co-localization. Finally, LC3C demonstrated high levels of co-localization, which significantly increased with the Y^86^ S mutant and decreased with the Y^247^ S and I^454^ T mutants (Figure 5 E, F; Suppl. Figure 4C). This fits to the observations of the co-immunoprecipitations. The Y^205^ S demonstrated an interesting dynamic for co-localization with LC3C, which decreased in cells not treated with Pyr-41.

**Figure 5.**
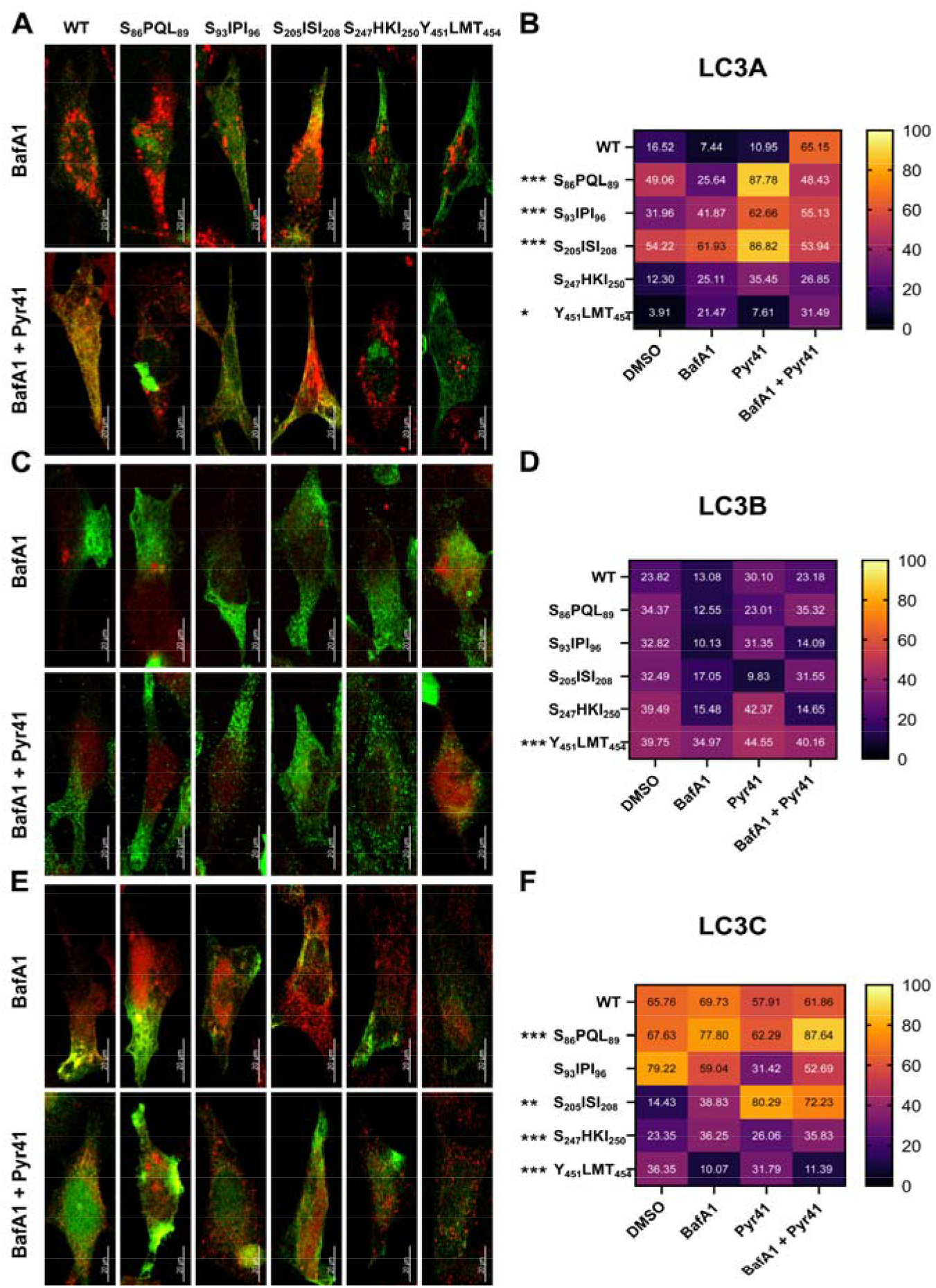
Co-localization of BAG3 and LC3s in single LIR mutants. (A) Representative images of *BAG3* ^-/-^ cells overexpressing single LIR mutants of BAG3 treated with Bafilomycin A1 (4 h, 2 μM) and PYR-41 (24 h, 12.5 μM). Magnification: 63x. Scale bar: 20 μm. BAG3 is shown in green, LC3A in red. (B) Heatmap comparing LC3A/BAG3 co-localizations among mutants and treatments. 2-way ANOVAs followed by a Benjamini Hochberg posthoc tests were used to determine statistical significance between the wildtype and the mutants over all treatment groups. ‘*’ indicates p<0.05 and ‘***’ p<0.001. (C) Representative images of *BAG3* ^-/-^ cells overexpressing single LIR mutants of BAG3 treated with Bafilomycin A1 (4 h, 2 μM) and PYR-41 (24 h, 12.5 μM). Magnification: 63x. Scale bar: 20 μm. BAG3 is shown in green, LC3B in red. (B) Heatmap comparing LC3B/BAG3 co-localizations among mutants and treatments. 2-way ANOVAs followed by a Benjamini Hochberg posthoc tests were used to determine statistical significance between the wildtype and the mutants over all treatment groups. ‘***’ indicates p<0.001. (A) Representative images of *BAG3* ^-/-^ cells overexpressing single LIR mutants of BAG3 treated with Bafilomycin A1 (4 h, 2 μM) and PYR-41 (24 h, 12.5 μM). Magnification: 63x. Scale bar: 20 μm. BAG3 is shown in green, LC3C in red. (B) Heatmap comparing LC3C/BAG3 co-localizations among mutants and treatments. 2-way ANOVAs followed by a Benjamini Hochberg posthoc tests were used to determine statistical significance. between the wildtype and the mutants over all treatment groups ‘**’ indicates p<0.01 and ‘***’ p<0.001.

Collectively, these data suggest that BAG3 may have multiple functional LIR motifs that may be accessed differently by LC3 orthologues. However, considering the effects of the Y^93^ S and I^454^ T these motifs are likely negligible. Both also demonstrated lowest affinities in the peptide array. The effects observed in the co-immunoprecipitations and co-localizations may be explained by other aspects. Due to the close proximity to Y^86^ PQL^89^, Y^93^ S may influence the other LIR activity. Finally, the I^454^ T mutant might cause BAG3 to become autophagosomal cargo. Thereby, co-localizations or co-immunoprecipitations with LC3A and LC3C may become less frequent.

## Discussion

We recently described the potential of BAG3 to serve as an autophagy receptor since BAG3 features a surprisingly strong conservation of putative LIR motifs within the craniotes (30). Therefore, we investigated *in vitro* and *in cellulo* whether these LIR motifs are functional. Our data not only indicate that BAG3 has three functional LIR motifs (Y^86^ PQL^89^, Y^205^ ISI^208^ and Y^247^ HKI^250^), but also suggest that each LC3 has a preferential interaction site (Figure 4, 5).

The concomitant strong conservation of the other predicted LIRs, Y^93^ IPI^96^ and Y^451^ LMI^454^, throughout vertebrate evolution may be explained by being part of the first IPV motif and being part of the BAG domain, respectively. Y^205^ ISI^208^ is equally a part of the second IPV motif. Interaction with a LC3 protein presumably influences binding of small HSPs to BAG3. Possibly, this might represent a potential switch for interactions with HSPB6/8 or LC3 proteins.

Previous data suggested that BAG3 may partake in autophagic processes independent of ubiquitin and may potentially deal also with non-ubiquitinated cargo (8). Therefore, we looked at the involvement of BAG3 in autophagy in presence of Pyr-41 and disabled E1 ubiquitin ligase activity. Interestingly, we still observed active, albeit weaker, recruitment of BAG3 (Figure 2 C). Mutation of all putative LIR motifs (ΔLIR-BAG3) caused a collapse of co-localization with LC3A and LC3C. Furthermore, the quantity of BAG3 in autophagic vesicles massively decreased in this case. In conclusion, the LIR motifs appear to be required for translocation of BAG3 into autophagosomes, even in cells without ubiquitination of cargo (figure 2).

Interestingly, BAG3 and its here generated mutants often behave in a similar way regardless of Pyr-41 presence. However, the S^205^ ISI^208^ mutant demonstrates varying behavior when challenged with Pyr-41 regarding colocalization with LC3C (Figure 5 E, F). In general, colocalization of LC3C with BAG3 is relative abundant. Yet, with the S^205^ ISI^208^ mutant this high level of colocalization only occured in presence of Pyr-41. This mutant also exhibited an increased affinity for LC3C in the immunoprecipitations (Figure 4D). Additionally, LC3C itself indicated a high affinity towards Y^247^ HKI^250^ in the peptide array (Figure 4B). Since only the peptide containing this motif at the free N-terminaus was detected, the absence of interaction for the other three peptides containing Y^247^ HKI^250^ might be due to a folding-related inaccessibility of the motif as well as to unspecific binding. Therefore, this result of the peptide array is hardly conclusive.

A previous mass spectrometry-based approach suggested a ubiquitination of K^249^ in BAG3 (37). Ubiquitination within a LIR motifs will most likely result in its dysfunction. This may offer the potential of an ubiquitination dependent regulation as autophagy receptor. The Pyr-41 dependent co-localization between LC3C and the S^205^ ISI^208^ mutant may therefore be a consequence of an activation of the Y^247^ HKI^250^.

Although LC3B is often considered the *bona fide* autophagy marker linking ubiquitin receptors with their cargo to the autophagososmal membrane, it is not strictly necessary for autophagosome formation or cargo recognition (35). Indeed, other LC3s may very well compensate for a loss of LC3B (35). Furthermore, the phagophore is not exclusively equipped with a single type of LC3. However, LC3B and LC3C appear to work at opposing ends of the autophagic spectrum (38, 39). Collectively, our findings regarding selectivity of a LIR-dependent interaction of BAG3 with LC3 proteins suggest a preference of LC3C and LC3A relative to LC3B. Similar effects have been described for other autophagy receptors (28). The co-localization analyses with the WT-BAG3 and the ΔLIR-BAG3 might suggest the possibility of a LC3B specific but LIR independent interaction. Whereas the co-localization with LC3A and LC3C significantly decreased with disruption of all putative LIR motifs, LC3B increasingly co-localized with BAG3 (Figure 3 B). Further support for this might be provided by the peptide array, which likewise suggested a non-LIR dependent binding solely for LC3B (Figure 4B). Therefore, the ability of BAG3 to interact with all three LC3s underlines a possible importance for its function as an autophagy receptor.

Protein turnover in general is associated with ubiquitination of lysine residues (40). This also includes specialized types of autophagy that turnover entire organelles like mitophagy (41). Although ubiquitin independent occurrences of selective autophagy have been discussed, much of its physiological purpose remains unclear (42). However, the rather apparent exclusiveness of LC3C interacting with Y^247^ HKI^250^, which may possibly be regulated by ubiquitination, may offer an interesting point of view where exactly a ubiquitination independent form of autophagy may come into play. LC3C is often associated with processes like xenophagy and aggrephagy (24, 43). In both cases, the potential cargo may be unable to be ubiquitinated due to structural inaccessibility.BAG3 has not been implicated with xenophagic processes so far but is a very well recognized actor in aggrephagic processes (11, 44, 45). In this case, BAG3 does not necessarily recognize its cargo through ubiquitin binding, which makes it a possible mediator of ubiquitin independent autophagy (46). Therefore, if the cell access to its ubiquitination machinery is impaired, BAG3 may still be able to drive autophagy where it is necessary.

## Data availability statement

All data are incorporated into the article and its online supplementary material.

## Acknowledgments

We thank Michael Plenikowski for the preparation of the graphical abstract.

## Author Contributions

HK, MB and CB conceptualized the study. HK and MB drafted the manuscript with input from all authors. HK, MB, DS and HN performed experiments.

## Funding

This work was supported by the Deutsche Forschungsgemeinschaft (DFG, German Research Foundation) Project-ID 259130777-SFB 1177.

## Conflict of Interest Statement

None of the authors has competing interests.

## Notes

### Competing Interest Statement

The authors have declared no competing interest.

## References

1. Chiang, H.-L., Terlecky, S. R., Plant, C. P., and Dice, J. F. (1989) A Role for a 70-Kilodalton Heat Shock Protein in Lysosomal Degradation of Intracellular Proteins. Science. 246, 382–385

2. Kirchner, P., Bourdenx, M., Madrigal-Matute, J., Tiano, S., Diaz, A., Bartholdy, B. A., Will, B., and Cuervo, A. M. (2019) Proteome-wide analysis of chaperone-mediated autophagy targeting motifs. PLOS Biol. 17, e3000301

3. Lüders, J., Demand, J., and Höhfeld, J. (2000) The Ubiquitin-related BAG-1 Provides a Link between the Molecular Chaperones Hsc70/Hsp70 and the Proteasome. J. Biol. Chem. 275, 4613–4617

4. Kim, Y. E., Hipp, M. S., Bracher, A., Hayer-Hartl, M., and Ulrich Hartl, F. (2013) Molecular Chaperone Functions in Protein Folding and Proteostasis. Annu. Rev. Biochem. 82, 323–355

5. Garrido, C., Paul, C., Seigneuric, R., and Kampinga, H. H. (2012) The small heat shock proteins family: The long forgotten chaperones. Int. J. Biochem. Cell Biol. 44, 1588–1592

6. Gamerdinger, M., Hajieva, P., Kaya, A. M., Wolfrum, U., Hartl, F. U., and Behl, C. (2009) Protein quality control during aging involves recruitment of the macroautophagy pathway by BAG3. EMBO J. 28, 889–901

7. Takayama, S., Xie, Z., and Reed, J. C. (1999) An evolutionarily conserved family of Hsp70/Hsc70 molecular chaperone regulators. J. Biol. Chem. 274, 781–786

8. Gamerdinger, M., Kaya, A. M., Wolfrum, U., Clement, A. M., and Behl, C. (2011) BAG3 mediates chaperone-based aggresome-targeting and selective autophagy of misfolded proteins. EMBO Rep. 12, 149–156

9. Meriin, A. B., Narayanan, A., Meng, L., Alexandrov, I., Varelas, X., Cissé, I. I., and Sherman, M. Y. (2018) Hsp70–Bag3 complex is a hub for proteotoxicity-induced signaling that controls protein aggregation. Proc. Natl. Acad. Sci. 115, E7043–E7052

10. Carra, S., Seguin, S. J., Lambert, H., and Landry, J. (2008) HspB8 chaperone activity toward poly(Q)-containing proteins depends on its association with Bag3, a stimulator of macroautophagy. J. Biol. Chem. 283, 1437–1444

11. Klimek, C., Kathage, B., Wördehoff, J., and Höhfeld, J. (2017) BAG3-mediated proteostasis at a glance. J. Cell Sci. 130, 2781–2788

12. Minoia, M., Boncoraglio, A., Vinet, J., Morelli, F. F., Brunsting, J. F., Poletti, A., Krom, S., Reits, E., Kampinga, H. H., and Carra, S. (2014) BAG3 induces the sequestration of proteasomal clients into cytoplasmic puncta. Autophagy. 10, 1603–1621

13. Korac, J., Schaeffer, V., Kovacevic, I., Clement, A. M., Jungblut, B., Behl, C., Terzic, J., and Dikic, I. (2013) Ubiquitin-independent function of optineurin in autophagic clearance of protein aggregates. J. Cell Sci. 126, 580–592

14. Bjørkøy, G., Lamark, T., Brech, A., Outzen, H., Perander, M., Øvervatn, A., Stenmark, H., and Johansen, T. (2005) p62/SQSTM1 forms protein aggregates degraded by autophagy and has a protective effect on huntingtin-induced cell death. J. Cell Biol. 171, 603–614

15. Pankiv, S., Clausen, T. H., Lamark, T., Brech, A., Bruun, J.-A., Outzen, H., Øvervatn, A., Bjørkøy, G., and Johansen, T. (2007) p62/SQSTM1 Binds Directly to Atg8/LC3 to Facilitate Degradation of Ubiquitinated Protein Aggregates by Autophagy. J. Biol. Chem. 282, 24131–24145

16. Ichimura, Y., Kirisako, T., Takao, T., Satomi, Y., Shimonishi, Y., Ishihara, N., Mizushima, N., Tanida, I., Kominami, E., Ohsumi, M., Noda, T., and Ohsumi, Y. (2000) A ubiquitin-like system mediates protein lipidation. Nature. 408, 488–492

17. Birgisdottir, Å. B., Lamark, T., and Johansen, T. (2013) The LIR motif - crucial for selective autophagy. J. Cell Sci. 126, 3237–3247

18. Johansen, T., and Lamark, T. (2020) Selective Autophagy: ATG8 Family Proteins, LIR Motifs and Cargo Receptors. J. Mol. Biol. 432, 80–103

19. Lv, M., Wang, C., Li, F., Peng, J., Wen, B., Gong, Q., Shi, Y., and Tang, Y. (2017) Structural insights into the recognition of phosphorylated FUNDC1 by LC3B in mitophagy. Protein Cell. 8, 25–38

20. Birgisdottir, Å. B., Mouilleron, S., Bhujabal, Z., Wirth, M., Sjøttem, E., Evjen, G., Zhang, W., Lee, R., O’Reilly, N., Tooze, S. A., Lamark, T., and Johansen, T. (2019) Members of the autophagy class III phosphatidylinositol 3-kinase complex I interact with GABARAP and GABARAPL1 via LIR motifs. Autophagy. 15, 1333–1355

21. Rogov, V. V., Suzuki, H., Marinković, M., Lang, V., Kato, R., Kawasaki, M., Buljubašić, M.,prung, M., Rogova, N., Wakatsuki, S., Hamacher-Brady, A., Dötsch, V., Dikic, I., Brady, N. R., and Novak, I. (2017) Phosphorylation of the mitochondrial autophagy receptor Nix enhances its interaction with LC3 proteins. Sci. Rep. 7, 1–12

22. Keown, J. R., Black, M. M., Ferron, A., Yap, M., Barnett, M. J., Pearce, F. G., Stoye, J. P., and Goldstone, D. C. (2018) A helical LC3-interacting region mediates the interaction between the retroviral restriction factor Trim5α and mammalian autophagy-related ATG8 proteins. J. Biol. Chem. 293, 18378–18386

23. Huber, J., Obata, M., Gruber, J., Akutsu, M., Löhr, F., Rogova, N., Güntert, P., Dikic, I., Kirkin, V., Komatsu, M., Dötsch, V., and Rogov, V. V. (2020) An atypical LIR motif within UBA5 (ubiquitin like modifier activating enzyme 5) interacts with GABARAP proteins and mediates membrane localization of UBA5. Autophagy. 16, 256–270

24. von Muhlinen, N., Akutsu, M., Ravenhill, B. J., Foeglein, Á., Bloor, S., Rutherford, T. J., Freund, S. M. V., Komander, D., and Randow, F. (2012) LC3C, Bound Selectively by a Noncanonical LIR Motif in NDP52, Is Required for Antibacterial Autophagy. Mol. Cell. 48, 329–342

25. Marshall, R. S., Hua, Z., Mali, S., McLoughlin, F., and Vierstra, R. D. (2019) ATG8-Binding UIM Proteins Define a New Class of Autophagy Adaptors and Receptors. Cell. 177, 766-781.e24

26. Wirth, M., Zhang, W., Razi, M., Nyoni, L., Joshi, D., O’Reilly, N., Johansen, T., Tooze, S. A., and Mouilleron, S. (2019) Molecular determinants regulating selective binding of autophagy adapters and receptors to ATG8 proteins. Nat. Commun. 10.1038/s41467-019-10059-6

27. Atkinson, J. M., Ye, Y., Gebru, M. T., Liu, Q., Zhou, S., Young, M. M., Takahashi, Y., Lin, Q., Tian, F., and Wang, H.-G. (2019) Time-resolved FRET and NMR analyses reveal selective binding of peptides containing the LC3-interacting region to ATG8 family proteins. J. Biol. Chem. 294, 14033– 14042

28. Sora, V., Kumar, M., Maiani, E., Lambrughi, M., Tiberti, M., and Papaleo, E. (2020) Structure and Dynamics in the ATG8 Family From Experimental to Computational Techniques. Front. Cell Dev. Biol. 8, 1–28

29. García-Mata, R., Bebök, Z., Sorscher, E. J., and Sztul, E. S. (1999) Characterization and Dynamics of Aggresome Formation by a Cytosolic Gfp-Chimera. J. Cell Biol. 146, 1239–1254

30. Baeken, M. W., and Behl, C. (2022) On the origin of BAG(3) and its consequences for an expansion of BAG3’s role in protein homeostasis. J. Cell. Biochem. 123, 102–114

31. Gossen, M., Freundlieb, S., Bender, G., Müller, G., Hillen, W., and Bujard, H. (1995) Transcriptional Activation by Tetracyclines in Mammalian Cells. Science. 268, 1766–1769

32. Ran, F. A., Hsu, P. D., Wright, J., Agarwala, V., Scott, D. A., and Zhang, F. (2013) Genome engineering using the CRISPR-Cas9 system. Nat. Protoc. 8, 2281–2308

33. Schmitt, D., Bozkurt, S., Henning-domres, P., Huesmann, H., Eimer, S., Bindila, L., Tascher, G., Münch, C., Behl, C., and Kern, A. (2022) Protein content and lipid profiling of isolated native autophagosomes. EMBO Rep. e53065

34. Yang, Y., Kitagaki, J., Dai, R.-M., Tsai, Y. C., Lorick, K. L., Ludwig, R. L., Pierre, S. A., Jensen, J. P., Davydov, I. V., Oberoi, P., Li, C.-C. H., Kenten, J. H., Beutler, J. A., Vousden, K. H., and Weissman, A. M. (2007) Inhibitors of Ubiquitin-Activating Enzyme (E1), a New Class of Potential Cancer Therapeutics. Cancer Res. 67, 9472–9481

35. Baeken, M. W., Weckmann, K., Diefenthäler, P., Schulte, J., Yusifli, K., Moosmann, B., Behl, C., and Hajieva, P. (2020) Novel Insights into the Cellular Localization and Regulation of the Autophagosomal Proteins LC3A, LC3B and LC3C. Cells. 9, 2315

36. Nomoto, A., Nishinami, S., and Shiraki, K. (2021) Solubility Parameters of Amino Acids on Liquid– Liquid Phase Separation and Aggregation of Proteins. Front. Cell Dev. Biol. 9, 1–7

37. Akimov, V., Barrio-Hernandez, I., Hansen, S. V. F., Hallenborg, P., Pedersen, A.-K., Bekker-Jensen, D. B., Puglia, M., Christensen, S. D. K., Vanselow, J. T., Nielsen, M. M., Kratchmarova, I., Kelstrup, C. D., Olsen, J. V., and Blagoev, B. (2018) UbiSite approach for comprehensive mapping of lysine and N-terminal ubiquitination sites. Nat. Struct. Mol. Biol. 25, 631–640

38. Lamark, T., and Johansen, T. (2021) Mechanisms of Selective Autophagy. Annu. Rev. Cell Dev. Biol. 37, 143–169

39. Varga, V. B., Keresztes, F., Sigmond, T., Vellai, T., and Kovács, T. (2022) The evolutionary and functional divergence of the Atg8 autophagy protein superfamily. Biol. Futur. 10.1007/s42977-022-00123-6

40. Chen, R.-H., Chen, Y.-H., and Huang, T.-Y. (2019) Ubiquitin-mediated regulation of autophagy. J. Biomed. Sci. 26, 80

41. Lazarou, M., Sliter, D. A., Kane, L. A., Sarraf, S. A., Wang, C., Burman, J. L., Sideris, D. P., Fogel, A. I., and Youle, R. J. (2015) The ubiquitin kinase PINK1 recruits autophagy receptors to induce mitophagy. Nature. 524, 309–314

42. Khaminets, A., Behl, C., and Dikic, I. (2016) Ubiquitin-Dependent And Independent Signals In Selective Autophagy. Trends Cell Biol. 26, 6–16

43. Liu, X., Li, Y., Wang, X., Xing, R., Liu, K., Gan, Q., Tang, C., Gao, Z., Jian, Y., Luo, S., Guo, W., and Yang, C. (2017) The BEACH-containing protein WDR81 coordinates p62 and LC3C to promote aggrephagy. J. Cell Biol. 216, 1301–1320

44. Hiebel, C., Stürner, E., Hoffmeister, M., Tascher, G., Schwarz, M., Nagel, H., Behrends, C., Münch, C., and Behl, C. (2020) BAG3 Proteomic Signature under Proteostasis Stress. Cells. 9, 2416

45. Lin, H., Deaton, C. A., and Johnson, G. V. W. (2022) Commentary: BAG3 as a Mediator of Endosome Function and Tau Clearance. Neuroscience. 10.1016/j.neuroscience.2022.05.002

46. Adriaenssens, E., Tedesco, B., Mediani, L., Asselbergh, B., Crippa, V., Antoniani, F., Carra, S., Poletti, A., and Timmerman, V. (2020) BAG3 Pro209 mutants associated with myopathy and neuropathy relocate chaperones of the CASA-complex to aggresomes. Sci. Rep. 10, 1–18

