## supplemental information for "Co-chaperone BAG3 directly target autophagic degradation via its LC3-interacting regions"

### Supplemental Figures

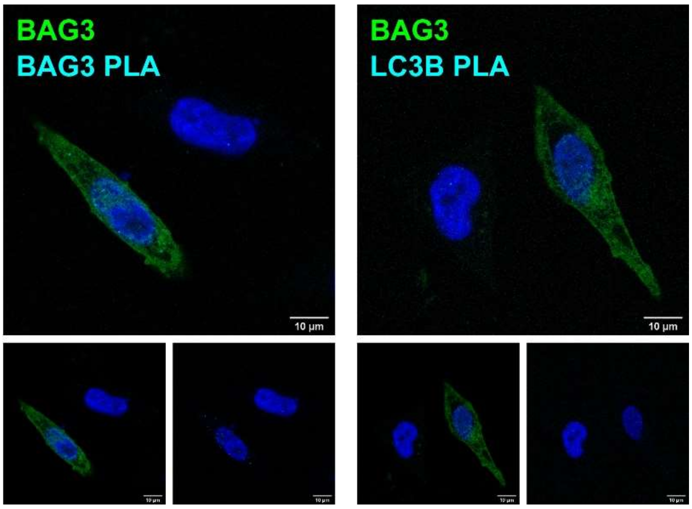


**Supplementary Figure 1:**

Negative controls to Figure 1. All PLAs were performed on *BAG3* $^{-/-}$ cells transiently transfected with human BAG3-eGFP and treated with MG-132 (10 h, 25 µM), Bafilomycin A1 (4 h, 2 µM) and PYR-41 (4 h, 12.5 µM). Representative images obtained from three biological replicates are displayed. Magnification: 100x. Scale bar: 10 µm.

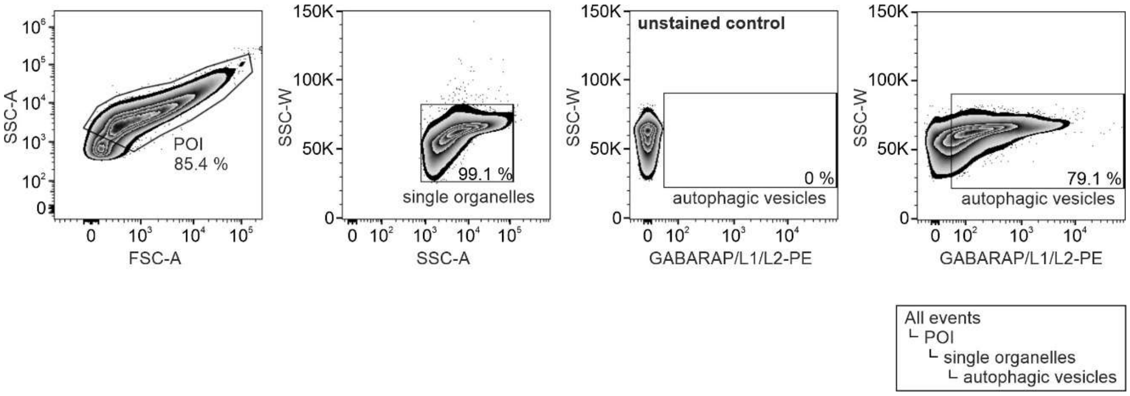


**Supplementary Figure 2:**

FACS-based isolation of autophagic vesicles. Representative images of the FACS gating strategy. The population of interest (POI) containing autophagic vesicles was first established using a FSC/SSC plot on a logarithmic scale, followed by a doublet discrimination using SSC-A/W. Autophagic vesicles were defined as PE-positive events (488 nm, BP 530/30) which were conducted according to the threshold given by an unstained negative control.

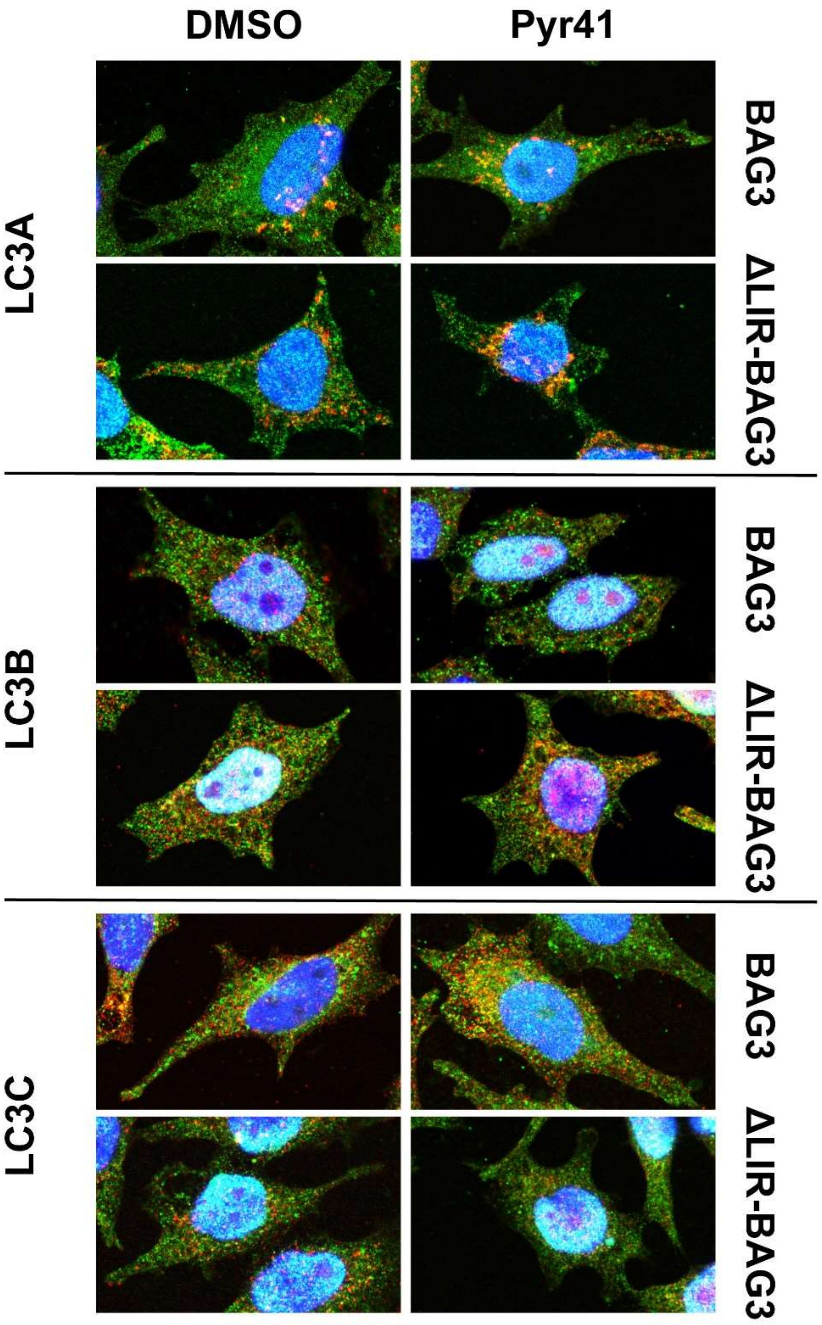


**Supplementary Figure 3:**

Minus Bafilomycin A1 controls to Figure 3. Representative images of *BAG3* $^{-/-}$ cells overexpressing WT-BAG3 or ΔLIR-BAG3 treated with DMSO and PYR-41 (24 h, 12.5 µM). Magnification: 100x. BAG3 is shown in green, the indicated LC3 in red.

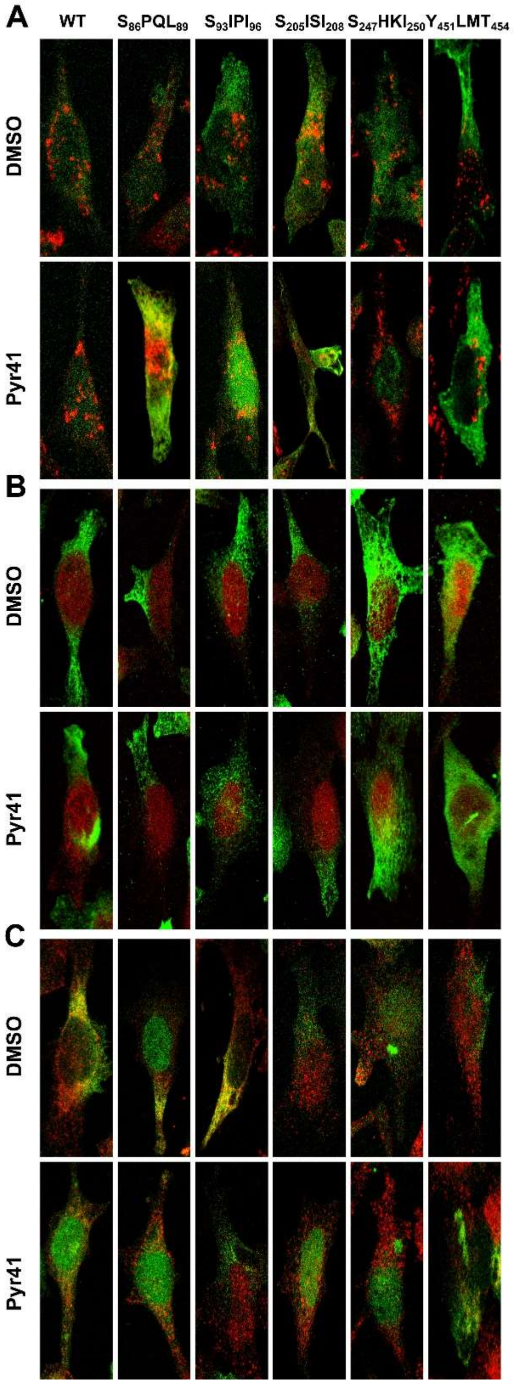


**Supplementary Figure 4:**

Minus Bafilomycin A1 controls to Figure 5. (A) Representative images of *BAG3*$^{-/-}$ cells overexpressing single LIR mutants of BAG3 treated with DMSO and PYR-41 (24 h, 12.5 µM). Magnification: 100x. Scale bar: 20 µm. BAG3 is shown in green, LC3A in red. (B) Representative images *BAG3*$^{-/-}$ cells overexpressing single LIR mutants of BAG3 treated with DMSO and PYR-41 (24 h, 12.5 µM). Magnification: 100x. Scale bar: 20 µm. BAG3 is shown in green, LC3B in red. (C) Representative images of *BAG3*$^{-/-}$ cells overexpressing single LIR mutants of BAG3 treated with DMSO and PYR-41 (24 h, 12.5 µM). Magnification: 100x. Scale bar: 20 µm. BAG3 is shown in green, LC3C in red.
